# Systematic assessment of the contribution of structural variants to inherited retinal diseases

**DOI:** 10.1101/2023.01.02.522522

**Authors:** Shu Wen, Meng Wang, Xinye Qian, Yumei Li, Keqing Wang, Jongsu Choi, Mark E. Pennesi, Paul Yang, Molly Marra, Robert K. Koenekoop, Irma Lopez, Anna Matynia, Michael Gorin, Ruifang Sui, Fengxia Yao, Kerry Goetz, Fernanda Belga Ottoni Porto, Rui Chen

**Affiliations:** Department of Molecular and Human Genetics, Baylor College of Medicine, Houston, Texas, United States; Human Genome Sequencing Center, Baylor College of Medicine, Houston, Texas, United States; Department of Ophthalmology, Casey Eye Institute, Oregon Health & Science University, Portland, OR, United States; McGill Ocular Genetics Laboratory and Centre, Department of Paediatric Surgery, Human Genetics, and Ophthalmology, McGill University Health Centre, Montreal, Quebec, Canada; Jules Stein Eye Institute, Los Angeles, California, United States; Ophthalmology, University of California Los Angeles David Geffen School of Medicine, Los Angeles, California, United States; Department of Ophthalmology, Peking Union Medical College Hospital, Peking Union Medical College, Chinese Academy of Medical Sciences, Beijing, China; Medical Research Center, State Key Laboratory of Complex Severe and Rare Diseases, Peking Union Medical College Hospital, Peking Union Medical College, Chinese Academy of Medical Sciences, China; Office of the Director, National Eye Institute/National Institutes of Health, Bethesda, MD, United States; INRET Clínica e Centro de Pesquisa, Belo Horizonte, Minas Gerais, Brazil; Department of Ophthalmology, Santa Casa de Misericórdia de Belo Horizonte, Belo Horizonte, Minas Gerais, Brazil; Centro Oftalmológico de Minas Gerais, Belo Horizonte, Minas Gerais, Brazil

**Author notes:** Corresponding author: Prof. Rui Chen,. Department of Molecular and Human Genetics, Baylor College of Medicine, Houston, Texas, United States.

## Abstract

Despite increasing success in determining genetic diagnosis for patients with inherited retinal diseases (IRDs), mutations in about 30% of the IRD cases remain unclear or unsettled after targeted gene panel or whole exome sequencing. In this study, we aimed to investigate the contributions of structural variants (SVs) to settling the molecular diagnosis of IRD with whole-genome sequencing (WGS). A cohort of 755 IRD patients whose pathogenic mutations remain undefined was subjected to WGS. Four SV calling algorithms including include MANTA, DELLY, LUMPY, and CNVnator were used to detect SVs throughout the genome. All SVs identified by any one of these four algorithms were included for further analysis. AnnotSV was used to annotate these SVs. SVs that overlap with known IRD-associated genes were examined with sequencing coverage, junction reads, and discordant read pairs. PCR followed by Sanger sequencing was used to further confirm the SVs and identify the breakpoints. Segregation of the candidate pathogenic alleles with the disease was performed when possible. In total, sixteen candidate pathogenic SVs were identified in sixteen families, including deletions and inversions, representing 2.1% of patients with previously unsolved IRDs. Autosomal dominant, autosomal recessive, and X-linked inheritance of disease-causing SVs were observed in 12 different genes. Among these, SVs in *CLN3, EYS, PRPF31* were found in multiple families. Our study suggests that the contribution of SVs detected by short-read WGS is about 0.25% of our IRD patient cohort and is significantly lower than that of single nucleotide changes and small insertions and deletions.

## INTRODUCTION

Structural variants (SVs) include genomic imbalance variants such as copy number variants (CNVs) due to deletions and duplications, and genomic balanced variants, including inversions and balanced translocations. The most studied type of SVs is CNV. Many methods, such as Multiplex Ligation-dependent Probe Amplification (MLPA), qPCR, Chromosome Microarray (CMA), targeted gene panel and WES, and WGS, have been used for CNV detection (1). For MLPA and qPCR, it is necessary to know the candidate CNV beforehand to design the primers or select the correct MLPA reagent. As the most frequently used technology in the detection of CNVs, CMA usually achieves a resolution as small as 5-10 kb (2, 3). However, because most CMA probes target the exons, when breakpoints are present in the introns or the intragenic regions, the identification of breakpoint location is much more difficult. Targeted-NGS-based CNV detection is becoming increasingly prevalent in recent years, however, they also pose similar weaknesses as CMA: the resolution is usually limited to two or more-consecutive exons, and the breakpoints cannot be mapped when they are located in non-targeted regions (4–6). Furthermore, it is difficult for CMA and targeted-NGS to detect balanced structural variants, such as inversions and translocations. Recently, with the rapid decline in prices of NGS, WGS is increasingly used for the detection of both SVs and single nucleotide variants (SNVs). Several studies have shown that WGS not only has excellent detection rate of SNVs, but also has a clear advantage in SV detection compared to other CNV detection methods (7–9). WGS is sensitive to CNVs of different sizes across the genome and can often map the breakpoints precisely regardless of their relative position to the exons. Another significant advantage of WGS-based SV analysis is that in addition to CNVs, it is possible to detect other types of SVs, such as an inversion and a translocation (7–9).

SVs have become increasingly recognized as a potential key genetic cause of human diseases due to improvement in new technologies, such as WGS technology (10, 11). Dozens of NGS based SV calling tools have been developed in the past couple of years, and their performances have been systematically compared (12). Previous studies have elucidated that although no tools are perfect, several tools such as MANTA (13), DELLY (14), LUMPY (15), CNVnator (16), have better overall performance (12). SVs of any size are potentially detectable by WGS. However, for short-read sequencing data, such as the ones provided by Illumina, SVs with lengths around 200– 500 bp are usually more difficult to detect due to the read-length limitation of the sequencing technology. Also, CNVs are easier to detect than balanced variants, because CNVs provided additional signals from gain or loss of read coverage in addition to discordant sequencing reads and paired reads.

Inherited retinal diseases (IRDs) are important causes of vision impairment that affect more than 2 million people worldwide. With the traditional genetic methods that focus on SNVs, such as targeted NGS, disease-causing SNVs can be detected in around 70% of all IRD patients. In addition, pathogenic SVs, mostly CNVs, have been reported in patients with IRDs (17–23). However, large scale systematic evaluation of SVs’ contribution to IRD has not been reported. In this study, application of Illumina short-read WGS to a large cohort of IRD patients with unsolved mutations after WES mutation screen allowed us to assess the contribution and variation spectrum of SVs among IRD patients.

## RESULTS

### Unsolved cohort of IRDs patients and initial SV discovery

To test the extent of SVs in IRD, we assembled a cohort of 755 previously unsolved patients (Fig. 1C). Retinitis pigmentosa (RP), Leber congenital amaurosis (LCA), cone-rod/cone dystrophy, *ABCA4-related* retinopathy (Stargardt disease), and Usher syndrome were the five most common diagnoses and accounted for more than 3/4 of the cohort. In short, these 755 IRD patients had undergone various types of molecular testing ranging from single gene tests to WES but remain unsettled.

**Fig. 1.**
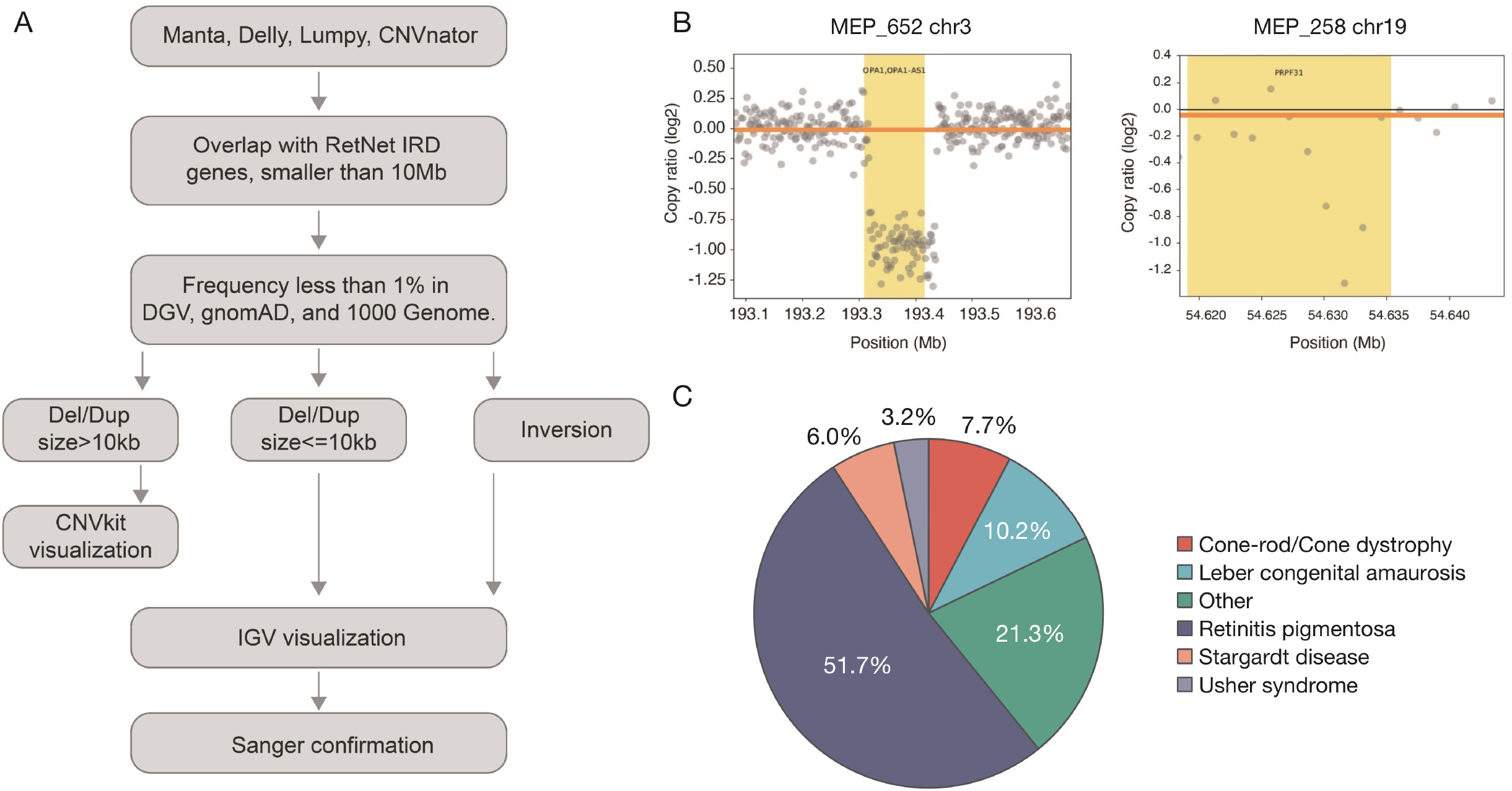
Analysis workflow and clinical information of the cohort. A. Informatics strategies used to filter and detect SVs. B. Larger CNVs are more easily to be called by CNVkit compared to smaller deletions. Larger deletions, using MEP_652’s 118 kb deletion (Panel A) as an example, contain dozens of bins, and are much more distinguishable compared to the baseline of smaller deletions, using MEP_258’s 5.2 kb deletion (Panel B) as an example. C. Clinical information of the 755 previously unsolved patients.

WGS was performed on these samples at a sequence depth of 30x. To identify candidate SVs at known IRD genes, a pipeline was developed as shown in Fig. 1A. To increase the sensitivity of SV calling, four software tools were used, including Manta, Delly, Lumpy, and CNVnator. As a result, a large number of SVs that overlapped with known IRD genes were identified, including 6535 deletion calls, 3924 duplication calls, and 48378 inversion calls (RetNet, https://sph.uth.edu/RetNet/). SVs were annotated to the RefSeq gene database and AnnotSV (24). A population frequency threshold of 1% was used in DGV, gnomAD, and 1000 Genome to filter out common SVs that occur too frequently to be the cause of rare IRDs (Fig. 1A). After filtering, 351 deletion calls, 239 duplication calls, and 304 inversion calls were kept for further analysis. Furthermore, SVs with high frequency in our internal database (>10%) were excluded as most of them were likely to be false positives. The remaining SVs that were larger than 10 kb were further visualized by CNVkit. Seven deletions and four duplications larger than 10 kb were consistent with the expected loss/gain of copy numbers in CNVkit were identified. In parallel, the bam files of the 19 deletions and eight duplications smaller than 10 kb, and all inversions were visualized in IGV. 1 deletion and 2 duplications smaller than 10 kb were consistent with the expected loss/gain of copy numbers in IGV. There was only one inversion that was consistent with the inversion reads in IGV and was then confirmed by Sanger sequencing of the breakpoints (data not shown). Finally, all SVs were combined with the SNV results and the clinical/pedigree information of each case for further pathogenicity analysis. As a result, a total of 16 highly confident deletions that were likely to be pathogenic were identified, including six large deletions greater than 10 kb and ten small deletions ranging from 966 to 9722 base pairs. No duplication or inversion was found to meet these criteria. All 16 deletions are further confirmed by PCR followed by Sanger sequencing to identify the breakpoints. Their details were summarized in Table 1.

**Table 1.**
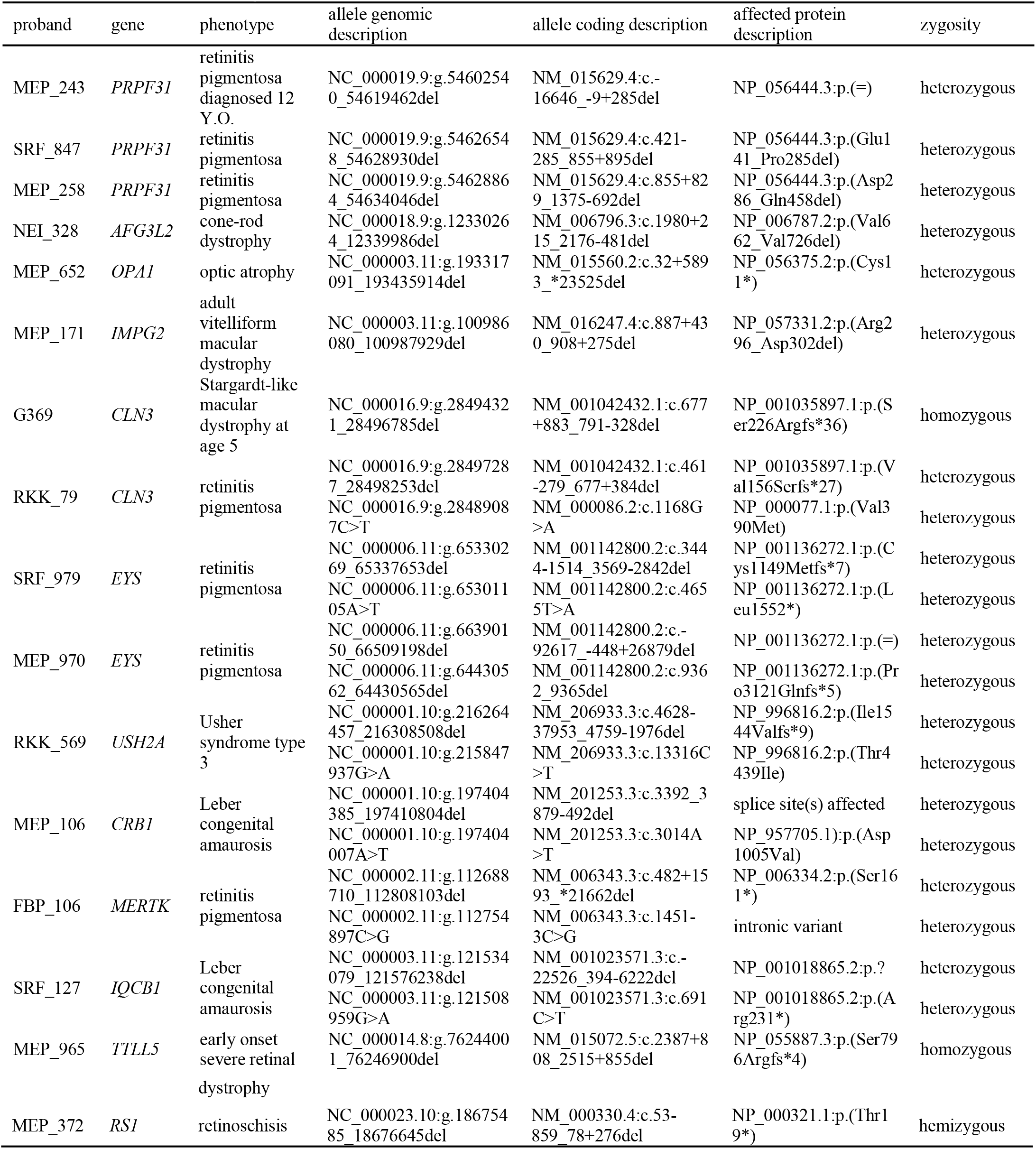
Summary of 16 probands carrying SVs in known IRD genes.

### Disease-associated SVs in Autosomal Dominant Genes

As shown in Table 1, a total of eight deletions have been found in genes that are associated with dominant IRDs, including *PRPF31, AFG3L2, OPA1*, and *IMPG2*. All but one variant is novel.

Among our patient cohort, three SV mutant alleles were identified in *PRPF31*, representing the most frequently mutated gene in this cohort. It has been shown that loss of function of one copy of *PRPF31* leads to RP due to haploinsufficiency (25). As shown in Table 1, all three alleles are novel and remove multiple coding exons of *PRPF31*. In addition, none of these three alleles have been observed in current population databases, such as gnomAD SV and DGV. All patients are diagnosed with RP, consistent with the molecular diagnosis. Specifically, MEP_243, who was diagnosed with RP at 12 years old (Fig. 2A), carries heterozygous c.-16646_-9+285 deletion that removes the first coding exon and the 16kb upstream sequences including the promoter and likely additional regulatory elements (Fig. 3A). Consistent with *PRPF31* as a dominant disease gene, the father of the proband is also affected. The proband started to notice decreased peripheral vision at 11. Her fundus images showed white spots of gliosis in both eyes. Her ERG showed unrecordable rod-dependent responses, as well as severely abnormal amplitudes and implicit times of the cone-dependent responses resulting in a pattern of rod-cone dysfunction (Fig. 2A). SRF_847 is a 23 year old female that carries a heterozygous c.421-285_855+895 deletion that removes exons 6-8, which is predicted to cause a frameshift (Fig. 3B). MEP_258 is a female patient diagnosed with RP at 28 years of age. Her daughter has similar symptoms, which support autosomal dominant inheritance. Her fundus showed RPE mottling with occasional bone spicules. Her ERG showed unrecordable rod-dependent responses, as well as severely abnormal amplitudes and abnormal implicit times of the cone-dependent responses (Fig. 2B). She carries an inframe heterozygous c.855+829_1375-692 deletion that removes exons 9-13 (Fig. 3C), which is more than 10% of the coding region, including the entire nuclear localization signal (26), a critical domain for *PRPF31* protein as part of the spliceosome complex. Furthermore, in ClinVar (27), three known pathogenic missense mutations have been mapped to exon 9-13, indicating the importance of this region for protein function. Taken together, all three *PRPF31* deletions are likely to be pathogenic mutations.

**Fig. 2.**
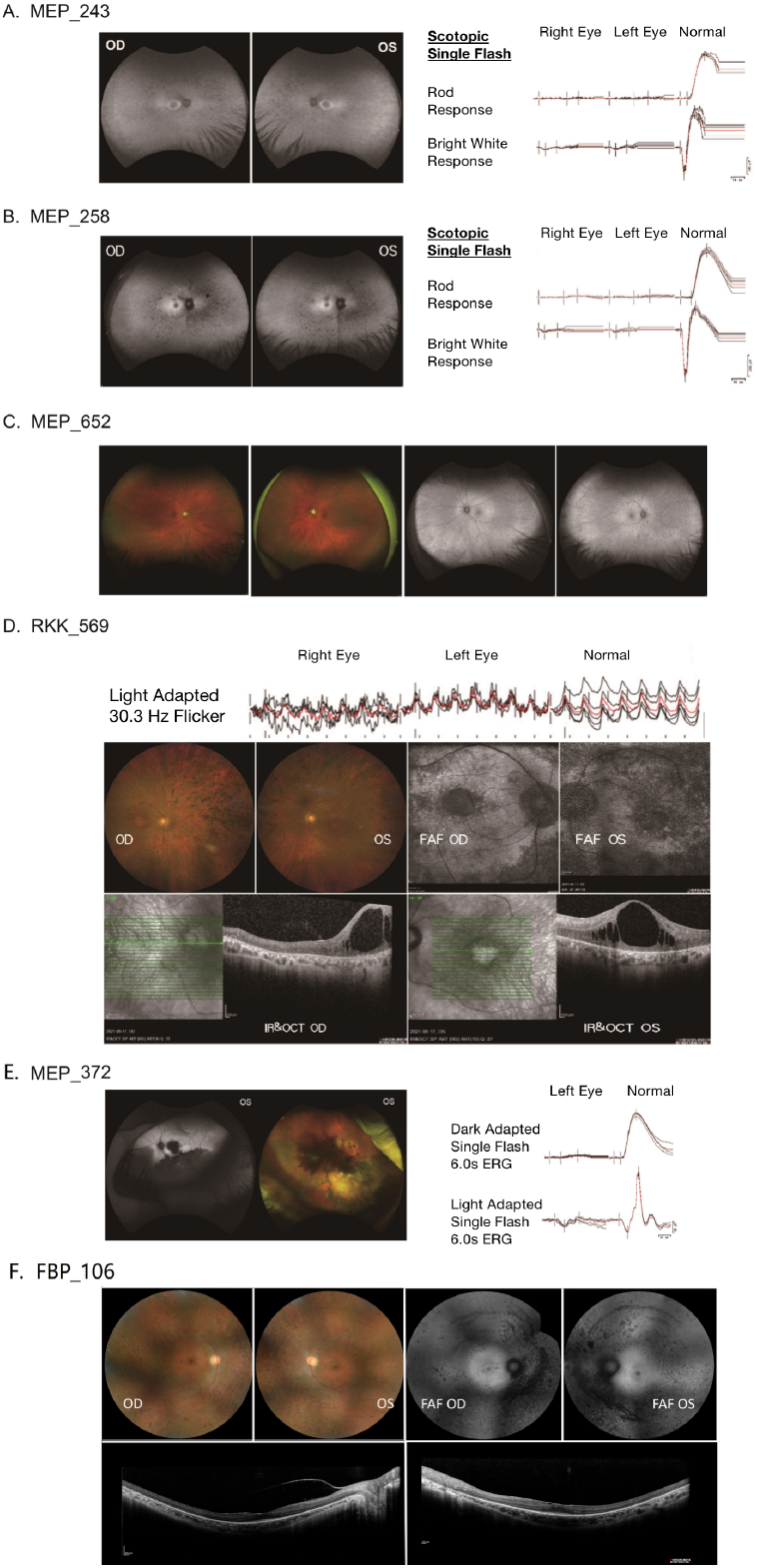
Clinical data supported the SVs’ deleterious effects based on known genotype-phenotype associations. A. Fundus autofluorescence and Electroretinography (ERG) of MEP_243. B. Fundus autofluorescence and ERG of MEP_258. C. Fundus autofluorescence of MEP_652. D. Fundus images and ERG of RKK_569. E. Fundus images and ERG of MEP_372. F. Fundus images and OCT of FBP_106.

**Fig. 3.**
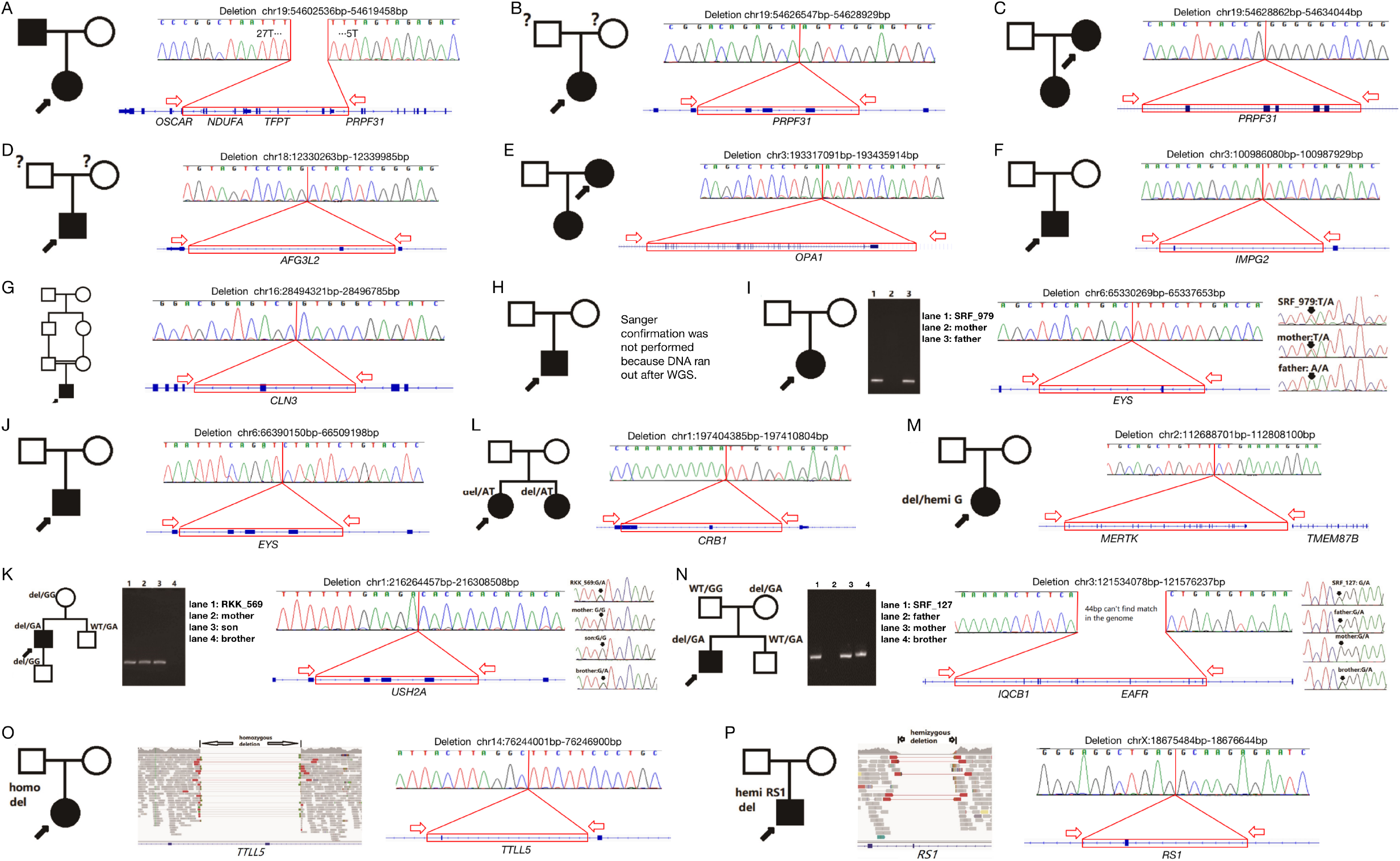
Probands’ pedigree, breakpoint PCR-electrophoresis, SV Sanger confirmation and segregation. A. A heterozygous *PRPF31* deletion which inherited from her father was detected in MEP_243 pedigree. B. A heterozygous deletion in *PRPF31* was detected in MEP_243 pedigree. C. A heterozygous *PRPF31* deletion which inherited to her daughter was detected in MEP_258 pedigree. D. A heterozygous deletion in *AFG3L2* was detected in NEI_328 pedigree. E. A heterozygous *OPA1* deletion which inherited to her daughter was detected in MEP_652 pedigree. F. A de novo heterozygous deletion in *IMPG2* was detected in MEP_171 pedigree. G. A *de novo* homozygous *CLN3* deletion was detected in a consanguineous family of G369. H. Compound heterozygous deletion variants of *CLN3* were detected in RKK_79 pedigree. Sanger confirmation was not performed because DNA ran out after WGS. I. Compound heterozygous variants of *EYS* were detected in SRF_979 pedigree. The compound heterozygous variants consisted of a deletion and a heterozygous variant of c.4655T>A (p.Leu1552*). J. Compound heterozygous variants of *EYS* were detected in MEP_970 pedigree. The compound heterozygous variants consisted of a deletion and a heterozygous variant of c.9362_9365del (p.Pro3121Glnfs*5). K. Compound heterozygous variants of *USH2A* were detected in RKK_569 pedigree. The compound heterozygous variants consisted of a deletion and a heterozygous variant of c. 13316C>T (p.Thr4439Ile). L. Compound heterozygous variants of *CRB1* were detected in MEP_106 and MEP_107’s pedigree. The compound heterozygous variants consisted of a heterozygous deletion and a heterozygous variant, c.3014A>T (p.Asp1005Val). M. A heterozygous *MERTK* deletion unmasks an overlapped hemizygous c.1451-3C>G intronic variant in *MERTK* gene was uncovered in FBP_106 pedigree. N. A heterozygous *IQCB1* deletion inherited from his asymptomatic father and a heterozygous c.691C>T(p.Arg231*) inherited from his asymptomatic mother were detected in SRF_127 pedigree. O. A *de novo* homozygous *TTLL5* deletion was identified in MEP_965 pedigree. P. A hemizygous *RS1* deletion was identified in MEP_372 pedigree.

Patient NEI_328 is a 49-year-old African American male that was diagnosed with cone-rod dystrophy (Fig. 2C). In NEI_328, an inframe heterozygous 9722-bp c.1980+215_2176-481 deletion that removes the penultimate exon of the *AFG3L2* was detected (Fig. 3D). Loss-of-function pathogenic variants in *AFG3L2* are known to cause AD optic atrophy 12 (28), which is consistent with the phenotype of our patient. This deletion was not seen in current population databases, such as gnomAD SV, DGV or literature. Furthermore, more than 10 pathogenic or likely pathogenic missense variants are reported mapped in the deleted region in ClinVar. Five of those variants (rs151344523, rs151344522, rs151344520, rs797045221, rs151344514) were also reported in Leiden Open Variation Database (LOVD) (29). Therefore, it is likely that this deletion is pathogenic.

Patient MEP_652 is a 56-year-old female diagnosed with optic atrophy (Fig. 2C). Her fundus images showed pallor of both optic nerves. Consistent with the clinical phenotype, a heterozygous c.32+5893_*23525 deletion that removes exon 2 to exon 29 (the last exon) of the *OPA1* was identified (Fig. 3E). Loss-of-function pathogenic variants in *OPA1* are known to cause AD optic atrophy 1 (30). This deletion is rare and is absent from population databases, such as gnomAD SV and DGV databases. In addition, the daughter of this individual, MEP_1016, who has similar optic atrophy as her mother, also carries this deletion, further supporting an autosomal dominant inheritance. Therefore, this *OPA1* deletion is considered pathogenic.

Patient MEP_171 is a 68-year-old male that has adult vitelliform macular dystrophy. He was diagnosed with macular degeneration at 54 years old. He currently has blurred vision and decreased night vision. His fundus images showed vitelliform foveal lesions with pigment clumping in a bull’s eye pattern (Fig. 2F). Consistent with the clinical phenotype, a heterozygous previously reported c.887+430_908+275 deletion is identified in *IMPG2*, heterozygous mutations in which lead to AD vitelliform macular dystrophy-5 (31). The deletion removes the entire exon 9 and was predicted to result in a frameshift (Fig. 3F). This allele is rare as it is absent from current population databases but has been reported in multiple vitelliform macular dystrophy patients (31). Therefore, it is likely the cause of the phenotype of MEP_171.

### Disease-associated SVs in autosomal recessive genes

A total of eight candidate causal pathogenic SV deletions were identified in genes associated with autosomal recessive IRDs, including 1 reported allele and 7 novel ones (Table 1).

Two deletions in *CLN3* have been found in the cohort. Loss-of-function pathogenic variants in *CLN3* are known to cause AR neuronal ceroid lipofuscinosis-3 (32), which often exhibits a severe cone-rod dystrophy. Patient G369 is a male who was initially diagnosed with Stargardt-like macular dystrophy at age 5. His parents were first cousins (Fig. 3G). There was no family history of eye disease. At 9 years old, he developed seizures and, later, muscle weakness. Biopsy of the conjunctiva supported the diagnosis of neuronal ceroid lipofuscinosis. The patient then lost speech and later died of pneumonia. A novel homozygous *CLN3* c.677+883_791-328 deletion that removes the entire exon 7 of *CLN3* was identified in the patient (Fig. 3G). This deletion results in a frameshift and was predicted to lead to NMD. This deletion is rare and has not been observed in population databases, such as gnomAD and DGV. Therefore, this *CLN3* deletion is likely pathogenic. The second proband, RKK_79, is a 66-year-old male who was diagnosed with RP (Fig. 3H). He is compound heterozygous for mutations in *CLN3*, including a previously reported frameshift deletion that removes the entire exon 9 and is predicted to cause NMD (33–36) and a c.1168G>A (p.Val390Met) missense mutation. The missense variant c.1168G>A (p.Val390Met) is rare with a population frequency of 3.58×10^-5^ and is consistently predicted to be deleterious by multiple programs such as REVEL (37) with a score of 0.98, a highly pathogenic score. Although both G369 and RKK_79 have been related to *CLN3*, RKK_79 has a much later age of onset and non-syndromic phenotype compared to G369. This is consistent with our and other groups’ finding (38–40) that non-syndromic *CLN3*-related RP cases are usually associated with at least one missense variant, whereas G369, the patient that carries the homozygous frameshift large deletion, has more severe neuronal ceroid lipofuscinosis-like symptoms, such as seizure and muscle weakness.

Mutations in *EYS*, which result in autosomal recessive retinitis pigmentosa 25, have been identified in two unrelated patients with RP (41). The first patient, SRF_979, is a 32-year-old female carrying compound heterozygous mutations, including a heterozygous frameshift c.3444-1514_3569-2842 deletion that removes the entire exon 22 and a heterozygous c.4655T>A (p.Leu1552*) nonsense mutation (Fig. 3I). Both mutations are predicted to cause NMD. Familial segregation analysis showed the father carries the deletion, and the mother carries the nonsense mutation (Fig. 3I). The second proband, MEP_970, carries a compound heterozygous deletion that removes 118kb of the *EYS* promoter and the entire exon 1 (Fig. 3J) and a heterozygous c.9362_9365del (p.Pro3121Glnfs*5) variant that was predicted to cause NMD (Fig. 3K). He is diagnosed with RP and is legally blind at 32 years old. He also has a sister with RP. Neither of their parents, or patient’s son, or brother were affected by IRDs. None of the four *EYS* variants has been observed in current population databases. Taken together, these *EYS* alleles are likely the cause of the phenotype of SRF_979 and MEP_970.

Patient RKK_569 is a male patient diagnosed with Usher syndrome type 3 (Fig. 2D). He has moderate to severe sensory-neural hearing loss from 0.25-8 kHz and needed bilateral hearing aid. His fundus images were consistent with retinitis pigmentosa. The patient carries a heterozygous c.4628-37953_4759-1976 deletion that removes entire exon 22 and is predicted to cause a frameshift deletion and NMD, and a known heterozygous pathogenic missense mutation c.13316C>T (p.Thr4439Ile) (42–44) in *USH2A* (Fig. 3K). *USH2A* is an autosomal recessive Usher syndrome gene, which is consistent with this patient’s phenotype. Neither the deletion nor the missense variant is seen in the population databases, such as gnomAD SV database or DGV. Family segregation analysis showed the mother and the son only carry the deletion, and the unaffected brother only carries the nonsense mutation (Fig. 3K). Taken together, these *USH2A* variants are likely the causes of the phenotype of this patient.

We identified a heterozygous deletion and a missense mutation in *CRB1* in two Caucasian sisters, MEP_106 and MEP_107. Both were diagnosed with LCA. Loss-of-function pathogenic variants in CRB1 are known to cause AR Leber congenital amaurosis 8 (45), which is consistent with the phenotype of our patient. The patient carries a heterozygous c.3392_3879-492 deletion, which removes part of exon 9 and the entire exon 10 of *CRB1* and results in a frameshift that was predicted to lead to NMD (Fig. 3L). This deletion has not been reported previously and is reported in the gnomAD SV database once. The patient also carries a known heterozygous c.3014A>T (p.Asp1005Val) pathogenic mutation (46, 47) in *CRB1*. Taken together, the deletion identified is considered to be pathogenic.

Patient FBP_106 is a 28-year-old female diagnosed with RP (Fig. 2G). She presented high bilateral myopia (−14, 0) and best-corrected visual acuity was 20/20. Examination showed thin retinal vessels, pigmentation in the periphery, temporal pallor of the optic discs, and flat electroretinographic responses. The patient carries a heterozygous c.482+1590_*21659 deletion that removes exon 3 through exon 19 (the last exon), which also unmasks an overlapping hemizygous c.1451-3C>G intronic variant in the *MERTK* (Fig. 3M). The c.1451-3C>G intronic variant was predicted by SpliceAI (48) to abolish the splice acceptor function with “high precision”. This likely leads to the skipping of exon 10, resulting in a shift in the open reading frame and NMD of the mRNA. Neither the deletion nor the splicing variant has been observed in population databases. Therefore, both variants are scored as pathogenic, and we believe they are the causes of the phenotype of this patient.

Patient SRF_127 is diagnosed with LCA and carries a heterozygous c.-22526_394-6222 deletion that removes more than 10kb of the promoter (overlaps with *EAF2*) and exon 1-5, and a novel heterozygous c.691C>T (p.Arg231*) stop gain in exon 11 of 15 exons that is predicted to cause NMD *in trans* in *IQCB1* (Fig. 3N). This patient also has intellectual disability and epilepsy. Besides, she has increased urea and creatinine, indicating abnormal kidney function. His brother also has LCA, intellectual disability and epilepsy. His kidney and liver functions are currently normal. Loss-of-function pathogenic variants in *IQCB1* are known to cause AR Senior-Loken syndrome 5 (49), which often has retinitis pigmentosa and LCA phenotypes. The deletion is rare as it has not been observed in population databases. The stop gain variant has a low population frequency of 0.01% in the gnomAD database (Fig. 3K). Therefore, both variants are novel pathogenic mutations in *IQCB1* and we believe they lead to the patient’s phenotype.

Patient MEP_965 is an Asian female with early onset severe retinal dystrophy, optic nerve edema, high myopia, and ptosis. In this patient, a homozygous frameshift c.2387+808_2515+855 deletion was detected in *TTLL5*, which removes the entire exon 24 and was predicted to cause NMD (Fig. 3O). Pathogenic variants in *TTLL5* are known to cause cone-rod dystrophy 19 (50), which is consistent with our patient. This deletion is not observed in population databases. Therefore, this deletion is considered a novel pathogenic mutation in *TTLL5*.

### Disease-associated SV in A X-Linked Gene

Patient MEP_372 is a male diagnosed with X-linked retinoschisis at age of 11. At his most recent visit at age 43, his visual acuity was no light perception for the right eye and 20/300 for the left eye (NLP OD and 20/300 OS). His right eye has had multiple surgeries and cannot be viewed with fundoscopy or tested by ERG. Slit lamp examination of the left eye revealed a 2+ nuclear sclerotic and 3+ posterior subcapsular cataract. His left eye has schisis with inner retinal holes inferiorly and temporally, with scarring medially. The vessels are attenuated and a cataract can be seen. His ERG of left eye showed decreased amplitudes to the DA 0.01 stimulus with prolonged timing and a negative waveform to the DA 6.0 stimulus. Cone driven responses were also reduced with a decreased b: a-wave ratio to the LA 3.0 stimulus and decreased amplitude and prolonged timing the LA 30Hz stimulus (Fig. 2E). Consistent with the clinical phenotype, a hemizygous frameshift c.53-859_78+276 deletion was detected in *RS1*, which removes the entire exon 2 of *RS1*. The predicted deletion was 26 bp in length, leading to NMD (Fig. 3P). This variant is absent in population databases. Although this allele is novel, exon 2 deletions have been reported in multiple patients with retinoschisis (51). Taken together, this deletion is considered as pathogenic.

## DISCUSSION

In this study, we performed a systematic investigation and identification of putative pathogenic SVs associated with IRD by analyzing WGS from 755 IRD patients whose mutations remain unknown after screening for mutations in coding region and splicing sites of known IRD genes. Following a set of stringent criteria, a list of 16 pathogenic and likely pathogenic SVs has been identified, representing one of the largest studies focusing on SVs in IRD patients to date.

### Most of the SVs identified in the study are novel

One striking observation from our study is that 88% (14/16) of pathogenic and likely pathogenic SVs identified in this study are novel. This is in contrast to what has been observed in previous cohort studies focusing on other types of variants, such as SNVs and small insertions/deletions. In those, the proportion of variants that are novel ranges from 30% to 50% depending on the ethnicity, with Caucasian the lowest and African the highest (22, 47, 52–54). This difference is likely due to a combination of factors, including the lack of SV studies and the rarity of SVs in general. Indeed, none except one of the 16 SVs identified in this study have been observed in the population databases, indicating these variants are exceedingly rare. In contrast, the carrier frequency of pathogenic SNVs is higher.

### Only Pathogenic and likely pathogenic deletions were identified in this study

During the study, we did not find any duplications, inversions, or other complex rearrangements that could explain patients’ phenotypes. A few duplications and inversions were detected and confirmed, but none of them was able to explain patients’ phenotypes (data not shown). Nine out of sixteen disease-causing deletions only involved one to two exons (Table 1). With targeted NGS and CMA, the CNVs with small size are more likely to be missed (6, 55, 56). Our data showed that small CNVs affecting less than three exons can account for a significant portion of CNVs, and WGS can detect many of them with its unique combination of coverage depth and breakpoint detection. On the other hand, we were not able to detect any duplications and/or inversions that can explain patients’ phenotypes. This is probably due to the limitation of our algorithm and sequencing technologies. For example, our detection rate of inversion could be artificially low. Taken together, in our cohort, pathogenic and likely pathogenic SVs are highly skewed to deletions.

### Only a small portion of unsolved IRD patients is due to SVs affecting the coding region of known IRD genes

The overall yield of this study is low with 2.1% (16/755) of the unsolved cases resolved. There are several potential reasons. First, detecting SVs from WGS remains challenging. To address this issue, we applied the four top performing SV calling algorithms, including MANTA (13), DELLY (14), LUMPY (15), and CNVnator (16). These algorithms consider both coverage depth and breakpoints, which can provide more information compared to algorithms that rely on coverage depth alone. As shown in Table 2, there is no single algorithm that can capture all 29 SVs that were later confirmed by either CNVkit or Sanger sequencing of the breakpoints (Table 2). We found Manta to be most sensitive, which detected 22/29 SVs. There were 9 SVs that would have been missed if Manta was not used because none of the other three algorithms detected these 9 SVs. On the contrary, CNVnator only detected 7/29 SVs, with no single SV that was only detected by CNVnator and not by any other algorithms (Table 2). However, despite using four algorithms to call SVs, some true SVs are likely missed by all callers. Second, we have been primarily focusing on SVs overlap with the coding region of the genes. As a result, SVs that only affect regulatory elements of the gene will be missed in our study. Third, due to the short read length, SVs in duplicate or highly repetitive regions of the genome are likely missed by current technology due to difficulties of accurate read mapping. One possible solution to this issue is to apply long-read NGS methods, such as Oxford Nanopore and PacBio. Furthermore, phasing information obtained from long reads will provide valuable information to determine if the two mutations are on the same or different chromosomes, a critical information for analyzing recessive IRD cases. It is also worth noting that these 16 patients are among a large cohort of 6,532 IRD patients, among which 5,768 are due to single base or small in/del mutations. Therefore, SVs in known IRD genes only contribute to 0.25% (16/6,532) of our cohort as the cause of the disease based on our current analysis.

**Table 2.**
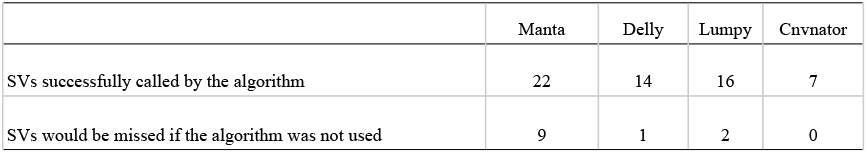
SVs being called and missed by each SV calling algorithm.

### SVs are more frequent in genes within Alu-rich regions

In our study, three different *PRPF31* deletions are found in three unrelated individuals. *PRPF31* has been reported by multiple groups to contain deletions/duplications that can cause retinitis pigmentosa (25, 57, 58) and was reported to account for 2.5% of autosomal dominant retinitis pigmentosa (59). One potential reason that *PRPF31* is enriched with SVs is that it is located in a region rich in repeat elements, especially Alu repeats (60). Alu-mediated rearrangements are one of the common causes of human disease-causing SVs (61). We also identified SVs in *CLN3* and *EYS* genes in each of two unrelated families that can explain their phenotypes. Interestingly, these two genes are also in Alu-rich regions (62, 63). As a comparison, the reported variants in ABCA4, one of the most common mutated gene in IRD patients that is not within Alu-rich region, are primarily SNVs and small in/dels (64). Therefore, thorough SV analysis is warranted particularly for genes mapped within Alu-rich region.

In summary, with 15/16 of pathogenic and likely pathogenic SVs identified in this study to be novel variants, it underscores the importance of investigating the contribution of SVs to disease burden since vast majority of the pathogenic SVs remain to be discovered. To improve our ability to identify pathogenic SVs, it is essential to apply improved short and long read sequencing technologies to further expand the effort to include both coding and noncoding part of the genome.

## METHODS

### Clinical diagnosis and patient recruitment

All probands discussed herein were clinically diagnosed with IRDs following a thorough ophthalmologic examination by a qualified inherited retinal disease specialist. This study was approved by the institutional ethics boards at each affiliated institution and adhered to the tenets of the declaration of Helsinki. Before blood collection, all probands and family members provided written informed consent for DNA analysis and received genetic counseling in accordance with established guidelines. DNA samples from patients and available relatives were obtained using the Qiagen or Chemagen blood genomic DNA extraction kit (Qiagen, Hilden, Germany). All samples were analyzed first with targeted IRD panel and samples whose variants remained unknown were then subjected to WES. Samples that lacked a confident molecular diagnosis after WES were assigned as “unsolved” for further WGS analysis.

### WGS, SV/SNV annotation, and filtering

WGS was performed for unsolved cases at a depth of about 30x coverage using the Illumina NovaSeq6000 platform at 2×150 bp. WGS data were processed using a pipeline modified from our previous WES data analysis pipeline (65)(Fig. 1A). Briefly, NGS sequencing reads were aligned to the human genome assembly (hg19) with BWA and variants are called using GATK 4 (66). SVs were identified using MANTA 1.2.2 (13), DELLY v0.7.8 (14), LUMPY 0.2.13 (15), and CNVnator v0.3 (16) with their default parameters. All SVs identified by any one of these four algorithms were included for further analysis. SVs are further filtered against DGV, gnomAD, and 1000 genome databases. SVs more frequent than 1% in any one of these databases were filtered out. Coding/intronic SNVs and small (<50bp) insertion-deletion variants (INDELs) were identified, filtered, and annotated as mentioned previously (67) for the following comprehensive SV/SNVs/INDELs variant analysis.

### SV analysis and validation

The candidate SVs were analyzed in three different groups: CNVs with sizes larger than 10kb, CNVs smaller than 10 kb, and inversions. The workflow we used for SV analysis of these three groups is as follows.

CNVs larger than 10 kb were visualized by CNVkit (68). We first randomly chose 50 WGS samples from our patient cohort and pooled them as the copy-neutral reference by using CNVkit, then compared each WGS data to this reference. Previously called CNVs and their flanking regions were input into CNVkit for visualizing the relative copy numbers compared to the pooled reference. A bin size of 1636 bp is used as determined by the CNVkit. Read coverage of each candidate CNVs that are larger than 10 kb are inspected (Fig. 1B). Integrative Genomics Viewer (IGV) is used to directly visualize for the local read coverage depth changes and paired-end reads that support the breakpoints. CNVs smaller than 10kbs and inversions were visualized and manually examined for breakpoint using IGV (Fig. 1B).

For CNVs/inversions that were supported by more than two junction reads in IGV, we used Sanger sequencing to confirm the breakpoints. If a clinically-relevant SNV was also called in this sample, Sanger confirmations in the sample were also performed. Segregation analysis in the families was also performed by Sanger sequencing.

## Acknowledgments

The authors would like to thank the patients and families for their enthusiastic participation. The DNA sample and data for participant NEI_328 described in this manuscript were obtained from the National Eye Institute – National Ophthalmic Genotyping and Phenotyping Network (eyeGENE^®^ - Protocol 06-EI-0236 which has been funded in part from the National Institutes of Health/National Eye Institute, under Contract No. HHS-N-260-2007-00001-C). We would like to thank the eyeGENE^®^ Research Group for their contribution.

## Author Contributors

All authors contributed to the study conception and design. Material preparation, data collection and analysis were performed by SW, MW, XYQ, YML, KQW and JSC. The first draft of the manuscript was written by SW and all authors commented on previous versions of the manuscript. All authors read and approved the final manuscript.

## Data Availability statement

The datasets generated during and/or analyzed during the current study are available from the corresponding author on reasonable request.

## Declarations

### Ethics approval and consent to participate

This study was performed in line with the principles of the Declaration of Helsinki. Approval was granted by the Ethics Committee of Baylor College of Medicine (Jan 21, 2021/IRB No. H-29697).

Consent for all patients or guardian is obtained by physicians who diagnosed and recruited patients in this study.

## REFERENCES

1. Zhao, M., Wang, Q., Wang, Q., Jia, P. and Zhao, Z. (2013) Computational tools for copy number variation (CNV) detection using next-generation sequencing data: features and perspectives. BMC Bioinformatics, 14 Suppl 11, S1.

2. Batzir, N.A., Shohat, M. and Maya, I. (2015) Chromosomal Microarray Analysis (CMA) a Clinical Diagnostic Tool in the Prenatal and Postnatal Settings. Pediatr Endocrinol Rev, 13, 448–454.

3. Hollenbeck, D., Williams, C.L., Drazba, K., Descartes, M., Korf, B.R., Rutledge, S.L., Lose, E.J., Robin, N.H., Carroll, A.J. and Mikhail, F.M. (2017) Clinical relevance of small copy-number variants in chromosomal microarray clinical testing. Genet Med, 19, 377–385.

4. Shen, R., Zhang, Z., Zhuang, Y., Yang, X. and Duan, L. (2021) A novel TUBG1 mutation with neurodevelopmental disorder caused by malformations of cortical development. Biomed Res Int, 2021, 6644274.

5. Strom, S.P., Hossain, W.A., Grigorian, M., Li, M., Fierro, J., Scaringe, W., Yen, H.Y., Teguh, M., Liu, J., Gao, H. et al. (2021) A Streamlined Approach to Prader-Willi and Angelman Syndrome Molecular Diagnostics. Front Genet, 12, 608889.

6. Moreno-Cabrera, J.M., del Valle, J., Castellanos, E., Feliubadaló, L., Pineda, M., Brunet, J., Serra, E., Capellà, G., Lázaro, C. and Gel, B. (2020) Evaluation of CNV detection tools for NGS panel data in genetic diagnostics. European Journal of Human Genetics, 28, 1645–1655.

7. Whitford, W., Lehnert, K., Snell, R.G. and Jacobsen, J.C. (2019) Evaluation of the performance of copy number variant prediction tools for the detection of deletions from whole genome sequencing data. J Biomed Inform, 94, 103174.

8. Lee, W.-P., Zhu, Q., Yang, X., Liu, S., Cerveira, E., Ryan, M., Mil-Homens, A., Bellfy, L., Ye, K., Zhang, C. et al. (2021) JAX-CNV: A whole genome sequencing-based algorithm for copy number detection at clinical grade level. medRxiv, in press., 2021.2003.2016.21252173.

9. Zhou, J., Yang, Z., Sun, J., Liu, L., Zhou, X., Liu, F., Xing, Y., Cui, S., Xiong, S., Liu, X. et al. (2021) Whole Genome Sequencing in the Evaluation of Fetal Structural Anomalies: A Parallel Test with Chromosomal Microarray Plus Whole Exome Sequencing. Genes (Basel), 12.

10. Kumaran, M., Cass, C.E., Graham, K., Mackey, J.R., Hubaux, R., Lam, W., Yasui, Y. and Damaraju, S. (2017) Germline copy number variations are associated with breast cancer risk and prognosis. Sci Rep, 7, 14621.

11. Takumi, T. and Tamada, K. (2018) CNV biology in neurodevelopmental disorders. Curr Opin Neurobiol, 48, 183–192.

12. Gabrielaite, M., Torp, M.H., Rasmussen, M.S., Andreu-Sánchez, S., Vieira, F.G., Pedersen, C.B., Kinalis, S., Madsen, M.B., Kodama, M., Demircan, G.S. et al. (2021) A Comparison of Tools for Copy-Number Variation Detection in Germline Whole Exome and Whole Genome Sequencing Data. Cancers (Basel), 13.

13. Chen, X., Schulz-Trieglaff, O., Shaw, R., Barnes, B., Schlesinger, F., Källberg, M., Cox, A.J., Kruglyak, S. and Saunders, C.T. (2016) Manta: rapid detection of structural variants and indels for germline and cancer sequencing applications. Bioinformatics, 32, 1220–1222.

14. Rausch, T., Zichner, T., Schlattl, A., Stütz, A.M., Benes, V. and Korbel, J.O. (2012) DELLY: structural variant discovery by integrated paired-end and split-read analysis. Bioinformatics, 28, i333–i339.

15. Layer, R.M., Chiang, C., Quinlan, A.R. and Hall, I.M. (2014) LUMPY: a probabilistic framework for structural variant discovery. Genome Biol, 15, R84.

16. Abyzov, A., Urban, A.E., Snyder, M. and Gerstein, M. (2011) CNVnator: an approach to discover, genotype, and characterize typical and atypical CNVs from family and population genome sequencing. Genome Res, 21, 974–984.

17. AlMoallem, B., Bauwens, M., Walraedt, S., Delbeke, P., De Zaeytijd, J., Kestelyn, P., Meire, F., Janssens, S., van Cauwenbergh, C., Verdin, H. et al. (2015) Novel FRMD7 Mutations and Genomic Rearrangement Expand the Molecular Pathogenesis of X-Linked Idiopathic Infantile Nystagmus. Invest Ophthalmol Vis Sci, 56, 1701–1710.

18. Coppieters, F., Todeschini, A.L., Fujimaki, T., Baert, A., De Bruyne, M., Van Cauwenbergh, C., Verdin, H., Bauwens, M., Ongenaert, M., Kondo, M. et al. (2015) Hidden Genetic Variation in LCA9-Associated Congenital Blindness Explained by 5’UTR Mutations and Copy-Number Variations of NMNAT1. Hum Mutat, 36, 1188–1196.

19. Eisenberger, T., Neuhaus, C., Khan, A.O., Decker, C., Preising, M.N., Friedburg, C., Bieg, A., Gliem, M., Charbel Issa, P., Holz, F.G. et al. (2013) Increasing the yield in targeted next-generation sequencing by implicating CNV analysis, non-coding exons and the overall variant load: the example of retinal dystrophies. PLoS One, 8, e78496.

20. Lindstrand, A., Davis, E.E., Carvalho, C.M., Pehlivan, D., Willer, J.R., Tsai, I.C., Ramanathan, S., Zuppan, C., Sabo, A., Muzny, D. et al. (2014) Recurrent CNVs and SNVs at the NPHP1 locus contribute pathogenic alleles to Bardet-Biedl syndrome. Am J Hum Genet, 94, 745–754.

21. Perez-Carro, R., Corton, M., Sánchez-Navarro, I., Zurita, O., Sanchez-Bolivar, N., Sánchez-Alcudia, R., Lelieveld, S.H., Aller, E., Lopez-Martinez, M.A., López-Molina, M.I. et al. (2016) Panel-based NGS Reveals Novel Pathogenic Mutations in Autosomal Recessive Retinitis Pigmentosa. Sci Rep, 6, 19531.

22. Pontikos, N., Arno, G., Jurkute, N., Schiff, E., Ba-Abbad, R., Malka, S., Gimenez, A., Georgiou, M., Wright, G., Armengol, M. et al. (2020) Genetic Basis of Inherited Retinal Disease in a Molecularly Characterized Cohort of More Than 3000 Families from the United Kingdom. Ophthalmology, 127, 1384–1394.

23. Shah, M., Shanks, M., Packham, E., Williams, J., Haysmoore, J., MacLaren, R.E., Németh, A.H., Clouston, P. and Downes, S.M. (2020) Next generation sequencing using phenotype-based panels for genetic testing in inherited retinal diseases. Ophthalmic Genet, 41, 331–337.

24. Geoffroy, V., Herenger, Y., Kress, A., Stoetzel, C., Piton, A., Dollfus, H. and Muller, J. (2018) AnnotSV: an integrated tool for structural variations annotation. Bioinformatics, 34, 3572–3574.

25. Abu-Safieh, L., Vithana, E.N., Mantel, I., Holder, G.E., Pelosini, L., Bird, A.C. and Bhattacharya, S.S. (2006) A large deletion in the adRP gene PRPF31: evidence that haploinsufficiency is the cause of disease. Mol Vis, 12, 384–388.

26. Deery, E.C., Vithana, E.N., Newbold, R.J., Gallon, V.A., Bhattacharya, S.S., Warren, M.J., Hunt, D.M. and Wilkie, S.E. (2002) Disease mechanism for retinitis pigmentosa (RP11) caused by mutations in the splicing factor gene PRPF31. Hum Mol Genet, 11, 3209–3219.

27. Landrum, M.J., Lee, J.M., Benson, M., Brown, G.R., Chao, C., Chitipiralla, S., Gu, B., Hart, J., Hoffman, D., Jang, W. et al. (2018) ClinVar: improving access to variant interpretations and supporting evidence. Nucleic Acids Res, 46, D1062–D1067.

28. Tulli, S., Del Bondio, A., Baderna, V., Mazza, D., Codazzi, F., Pierson, T.M., Ambrosi, A., Nolte, D., Goizet, C., Toro, C. et al. (2019) Pathogenic variants in the AFG3L2 proteolytic domain cause SCA28 through haploinsufficiency and proteostatic stress-driven OMA1 activation. J Med Genet, 56, 499–511.

29. Fokkema, I.F., Taschner, P.E., Schaafsma, G.C., Celli, J., Laros, J.F. and den Dunnen, J.T. (2011) LOVD v.2.0: the next generation in gene variant databases. Hum Mutat, 32, 557–563.

30. Marchbank, N.J., Craig, J.E., Leek, J.P., Toohey, M., Churchill, A.J., Markham, A.F., Mackey, D.A., Toomes, C. and Inglehearn, C.F. (2002) Deletion of the OPA1 gene in a dominant optic atrophy family: evidence that haploinsufficiency is the cause of disease. J Med Genet, 39, e47.

31. Bandah-Rozenfeld, D., Collin, R.W., Banin, E., van den Born, L.I., Coene, K.L., Siemiatkowska, A.M., Zelinger, L., Khan, M.I., Lefeber, D.J., Erdinest, I. et al. (2010) Mutations in IMPG2, encoding interphotoreceptor matrix proteoglycan 2, cause autosomal-recessive retinitis pigmentosa. Am J Hum Genet, 87, 199–208.

32. Munroe, P.B., Mitchison, H.M., O’Rawe, A.M., Anderson, J.W., Boustany, R.M., Lerner, T.J., Taschner, P.E., de Vos, N., Breuning, M.H., Gardiner, R.M. et al. (1997) Spectrum of mutations in the Batten disease gene, CLN3. Am J Hum Genet, 61, 310–316.

33. Järvelä, I., Mitchison, H.M., Munroe, P.B., O’Rawe, A.M., Mole, S.E. and Syvänen, A.C. (1996) Rapid diagnostic test for the major mutation underlying Batten disease. J Med Genet, 33, 1041–1042.

34. Järvelä, I., Autti, T., Lamminranta, S., Aberg, L., Raininko, R. and Santavuori, P. (1997) Clinical and magnetic resonance imaging findings in Batten disease: analysis of the major mutation (1.02-kb deletion). Ann Neurol, 42, 799–802.

35. Kitzmüller, C., Haines, R.L., Codlin, S., Cutler, D.F. and Mole, S.E. (2008) A function retained by the common mutant CLN3 protein is responsible for the late onset of juvenile neuronal ceroid lipofuscinosis. Hum Mol Genet, 17, 303–312.

36. (1995) Isolation of a novel gene underlying Batten disease, CLN3. The International Batten Disease Consortium. Cell, 82, 949–957.

37. Ioannidis, N.M., Rothstein, J.H., Pejaver, V., Middha, S., McDonnell, S.K., Baheti, S., Musolf, A., Li, Q., Holzinger, E., Karyadi, D. et al. (2016) REVEL: An Ensemble Method for Predicting the Pathogenicity of Rare Missense Variants. Am J Hum Genet, 99, 877–885.

38. Chen, F.K., Zhang, X., Eintracht, J., Zhang, D., Arunachalam, S., Thompson, J.A., Chelva, E., Mallon, D., Chen, S.C., McLaren, T. et al. (2019) Clinical and molecular characterization of non-syndromic retinal dystrophy due to c.175G>A mutation in ceroid lipofuscinosis neuronal 3 (CLN3). Doc Ophthalmol, 138, 55–70.

39. Ku, C.A., Hull, S., Arno, G., Vincent, A., Carss, K., Kayton, R., Weeks, D., Anderson, G.W., Geraets, R., Parker, C. et al. (2017) Detailed Clinical Phenotype and Molecular Genetic Findings in CLN3-Associated Isolated Retinal Degeneration. JAMA Ophthalmol, 135, 749–760.

40. Wang, F., Wang, H., Tuan, H.F., Nguyen, D.H., Sun, V., Keser, V., Bowne, S.J., Sullivan, L.S., Luo, H., Zhao, L. et al. (2014) Next generation sequencing-based molecular diagnosis of retinitis pigmentosa: identification of a novel genotype-phenotype correlation and clinical refinements. Hum Genet, 133, 331–345.

41. Abd El-Aziz, M.M., Barragan, I., O’Driscoll, C.A., Goodstadt, L., Prigmore, E., Borrego, S., Mena, M., Pieras, J.I., El-Ashry, M.F., Safieh, L.A. et al. (2008) EYS, encoding an ortholog of Drosophila spacemaker, is mutated in autosomal recessive retinitis pigmentosa. Nat Genet, 40, 1285–1287.

42. Baux, D., Blanchet, C., Hamel, C., Meunier, I., Larrieu, L., Faugère, V., Vaché, C., Castorina, P., Puech, B., Bonneau, D. et al. (2014) Enrichment of LOVD-USHbases with 152 USH2A genotypes defines an extensive mutational spectrum and highlights missense hotspots. Hum Mutat, 35, 1179–1186.

43. Le Quesne Stabej, P., Saihan, Z., Rangesh, N., Steele-Stallard, H.B., Ambrose, J., Coffey, A., Emmerson, J., Haralambous, E., Hughes, Y., Steel, K.P. et al. (2012) Comprehensive sequence analysis of nine Usher syndrome genes in the UK National Collaborative Usher Study. J Med Genet, 49, 27–36.

44. Lenassi, E., Robson, A.G., Luxon, L.M., Bitner-Glindzicz, M. and Webster, A.R. (2015) Clinical heterogeneity in a family with mutations in USH2A. JAMA Ophthalmol, 133, 352–355.

45. Lotery, A.J., Jacobson, S.G., Fishman, G.A., Weleber, R.G., Fulton, A.B., Namperumalsamy, P., Héon, E., Levin, A.V., Grover, S., Rosenow, J.R. et al. (2001) Mutations in the CRB1 gene cause Leber congenital amaurosis. Arch Ophthalmol, 119, 415–420.

46. Corton, M., Tatu, S.D., Avila-Fernandez, A., Vallespín, E., Tapias, I., Cantalapiedra, D., Blanco-Kelly, F., Riveiro-Alvarez, R., Bernal, S., García-Sandoval, B. et al. (2013) High frequency of CRB1 mutations as cause of Early-Onset Retinal Dystrophies in the Spanish population. Orphanet J Rare Dis, 8, 20.

47. Stone, E.M., Andorf, J.L., Whitmore, S.S., DeLuca, A.P., Giacalone, J.C., Streb, L.M., Braun, T.A., Mullins, R.F., Scheetz, T.E., Sheffield, V.C. et al. (2017) Clinically Focused Molecular Investigation of 1000 Consecutive Families with Inherited Retinal Disease. Ophthalmology, 124, 1314–1331.

48. Jaganathan, K., Kyriazopoulou Panagiotopoulou, S., McRae, J.F., Darbandi, S.F., Knowles, D., Li, Y.I., Kosmicki, J.A., Arbelaez, J., Cui, W., Schwartz, G.B. et al. (2019) Predicting Splicing from Primary Sequence with Deep Learning. Cell, 176, 535–548.e524.

49. Otto, E.A., Loeys, B., Khanna, H., Hellemans, J., Sudbrak, R., Fan, S., Muerb, U., O’Toole, J.F., Helou, J., Attanasio, M. et al. (2005) Nephrocystin-5, a ciliary IQ domain protein, is mutated in Senior-Loken syndrome and interacts with RPGR and calmodulin. Nat Genet, 37, 282–288.

50. Sergouniotis, P.I., Chakarova, C., Murphy, C., Becker, M., Lenassi, E., Arno, G., Lek, M., MacArthur, D.G., Bhattacharya, S.S., Moore, A.T. et al. (2014) Biallelic variants in TTLL5, encoding a tubulin glutamylase, cause retinal dystrophy. Am J Hum Genet, 94, 760–769.

51. Sauer, C.G., Gehrig, A., Warneke-Wittstock, R., Marquardt, A., Ewing, C.C., Gibson, A., Lorenz, B., Jurklies, B. and Weber, B.H. (1997) Positional cloning of the gene associated with X-linked juvenile retinoschisis. Nat Genet, 17, 164–170.

52. Weisschuh, N., Obermaier, C.D., Battke, F., Bernd, A., Kuehlewein, L., Nasser, F., Zobor, D., Zrenner, E., Weber, E., Wissinger, B. et al. (2020) Genetic architecture of inherited retinal degeneration in Germany: A large cohort study from a single diagnostic center over a 9-year period. Human Mutation, 41, 1514–1527.

53. Weisschuh, N., Mayer, A.K., Strom, T.M., Kohl, S., Glöckle, N., Schubach, M., Andreasson, S., Bernd, A., Birch, D.G., Hamel, C.P. et al. (2016) Mutation Detection in Patients with Retinal Dystrophies Using Targeted Next Generation Sequencing. PLoS One, 11, e0145951.

54. Chen, T.-C., Huang, D.-S., Lin, C.-W., Yang, C.-H., Yang, C.-M., Wang, V.Y., Lin, J.-W., Luo, A.C., Hu, F.-R. and Chen, P.-L. (2021) Genetic characteristics and epidemiology of inherited retinal degeneration in Taiwan. npj Genomic Medicine, 6, 16.

55. Zare, F., Dow, M., Monteleone, N., Hosny, A. and Nabavi, S. (2017) An evaluation of copy number variation detection tools for cancer using whole exome sequencing data. BMC Bioinformatics, 18, 286.

56. Singh, A.K., Olsen, M.F., Lavik, L.A.S., Vold, T., Drabløs, F. and Sjursen, W. (2021) Detecting copy number variation in next generation sequencing data from diagnostic gene panels. BMC Medical Genomics, 14, 214.

57. Villanueva, A., Willer, J.R., Bryois, J., Dermitzakis, E.T., Katsanis, N. and Davis, E.E. (2014) Whole exome sequencing of a dominant retinitis pigmentosa family identifies a novel deletion in PRPF31. Invest Ophthalmol Vis Sci, 55, 2121–2129.

58. Wang, L., Ribaudo, M., Zhao, K., Yu, N., Chen, Q., Sun, Q., Wang, L. and Wang, Q. (2003) Novel deletion in the pre-mRNA splicing gene PRPF31 causes autosomal dominant retinitis pigmentosa in a large Chinese family. Am J Med Genet A, 121a, 235–239.

59. Sullivan, L.S., Bowne, S.J., Seaman, C.R., Blanton, S.H., Lewis, R.A., Heckenlively, J.R., Birch, D.G., Hughbanks-Wheaton, D. and Daiger, S.P. (2006) Genomic rearrangements of the PRPF31 gene account for 2.5% of autosomal dominant retinitis pigmentosa. Invest Ophthalmol Vis Sci, 47, 4579–4588.

60. Xiao, X., Cao, Y., Zhang, Z., Xu, Y., Zheng, Y., Chen, L.J., Pang, C.P. and Chen, H. (2017) Novel Mutations in PRPF31 Causing Retinitis Pigmentosa Identified Using Whole-Exome Sequencing. Invest Ophthalmol Vis Sci, 58, 6342–6350.

61. Song, X., Beck, C.R., Du, R., Campbell, I.M., Coban-Akdemir, Z., Gu, S., Breman, A.M., Stankiewicz, P., Ira, G., Shaw, C.A. et al. (2018) Predicting human genes susceptible to genomic instability associated with Alu/Alu-mediated rearrangements. Genome Res, 28, 1228–1242.

62. Hehir-Kwa, J.Y., Marschall, T., Kloosterman, W.P., Francioli, L.C., Baaijens, J.A., Dijkstra, L.J., Abdellaoui, A., Koval, V., Thung, D.T., Wardenaar, R. et al. (2016) A high-quality human reference panel reveals the complexity and distribution of genomic structural variants. Nat Commun, 7, 12989.

63. Mitchison, H.M., Taschner, P.E., Kremmidiotis, G., Callen, D.F., Doggett, N.A., Lerner, T.J., Janes, R.B., Wallace, B.A., Munroe, P.B., O’Rawe, A.M. et al. (1997) Structure of the CLN3 gene and predicted structure, location and function of CLN3 protein. Neuropediatrics, 28, 12–14.

64. Cremers, F.P.M., Lee, W., Collin, R.W.J. and Allikmets, R. (2020) Clinical spectrum, genetic complexity and therapeutic approaches for retinal disease caused by ABCA4 mutations. Prog Retin Eye Res, 79, 100861.

65. Soens, Z.T., Li, Y., Zhao, L., Eblimit, A., Dharmat, R., Li, Y., Chen, Y., Naqeeb, M., Fajardo, N., Lopez, I. et al. (2016) Hypomorphic mutations identified in the candidate Leber congenital amaurosis gene CLUAP1. Genet Med, 18, 1044–1051.

66. McKenna, A., Hanna, M., Banks, E., Sivachenko, A., Cibulskis, K., Kernytsky, A., Garimella, K., Altshuler, D., Gabriel, S., Daly, M. et al. (2010) The Genome Analysis Toolkit: a MapReduce framework for analyzing next-generation DNA sequencing data. Genome Res, 20, 1297–1303.

67. Qian, X., Wang, J., Wang, M., Igelman, A.D., Jones, K.D., Li, Y., Wang, K., Goetz, K.E., Birch, D.G., Yang, P. et al. (2021) Identification of Deep-Intronic Splice Mutations in a Large Cohort of Patients With Inherited Retinal Diseases. Front Genet, 12, 647400.

68. Talevich, E., Shain, A.H., Botton, T. and Bastian, B.C. (2016) CNVkit: Genome-Wide Copy Number Detection and Visualization from Targeted DNA Sequencing. PLoS Comput Biol, 12, e1004873.

